# Stochastic Emergence of Irregular Infection Fronts in Motile Bacteria-Phage Populations

**DOI:** 10.1101/2025.10.29.685271

**Authors:** Laura Bergamaschi, Namiko Mitarai

## Abstract

Interactions between bacteriophages and motile bacteria can produce irregular spatial patterns. Here we show that these irregularities arise from stochastic infection dynamics at the single-cell level. We develop a discrete, stochastic model of phage–bacteria co-propagation, which represents bacteria and phage numbers as non-negative integers on a two-dimensional lattice, while nutrients and attractants are treated as real-valued fields. Stochastic rules govern bacterial growth, chemotactic movement, infection, and lysis, allowing spatial heterogeneity to emerge in agreement with experimentally observed asymmetric patterns. Simulations reveal that rare events, in which an infected bacterium migrates ahead of the front before lysis, locally seed new infection centers. The resulting front roughness is controlled by the product of burst size and adsorption rate, and is suppressed when the effective population size per lattice site increases or the variability of latent period decreases. These results link microscopic stochasticity to emergent spatial structure in phage–bacteria populations.

## 1 Introduction

Bacteriophages play a central role in microbial ecosystems [1], from the human gut to oceans and soil [2]. While phage–bacteria interactions have been extensively studied in well-mixed liquid cultures [3, 4], spatial structure and bacterial motility can fundamentally alter infection outcomes, promoting coexistence and diversification [5, 6, 7, 8, 9, 10, 11, 12, 13, 14, 15]. In structured environments (such as semi-solid agar), motile bacteria create their own nutrient gradients and chemotactically migrate along them [16, 17, 18, 19]. Traveling-wave solutions combining growth and chemotaxis have been analytically characterized [20], providing theoretical context for bacterial front propagation. These expanding fronts often outpace the spread of phages, introducing a new regime in which non-motile phages must essentially ‘hitchhike’ by infecting moving bacteria to avoid being left behind [21, 22, 23, 24]. Experimental studies of such co-propagating phage–bacteria populations have revealed complex spatiotemporal dynamics – for example, the formation of distinct bacterial and phage spread fronts, and lysis zones where bacteria are cleared by phage [21, 22, 23, 24]. Notably, several features of these relatively smooth and symmetric patterns have been captured by traditional continuous models using deterministic partial differential equations (PDEs) [21, 22, 23, 24].

However, experiments also suggest that chance events and variability can upset these smooth patterns. In the study by Ping et al. (2020), co-cultures of chemotactic *E. coli* and bacteriophage P1_vir_ in swimming agar sometimes developed irregular, rough-edged lysis patterns instead of smooth expanding rings [21]. These rough “killing fronts” occurred under intermediate phage concentrations and adsorption rates – conditions near a transition between phage falling behind vs. overtaking the bacteria. Such irregularities highlight the potential importance of randomness. Even without large initial perturbations, rare events can arise – for instance, a single bacterium infected at the leading edge might swim ahead of the pack and then lyse, seeding a new pocket of phages farther out. This one infection can lead to a burst of local phage that initiates a new infection focus, distorting the overall pattern. In other words, an initially minor, random event can be amplified by feedback into a macroscale asymmetry. However, such effects are inherently difficult to capture with PDE-based models. While deterministic PDEs can exhibit irregular fronts through intrinsic instabilities, such as viscous fingering in Laplacian growth models [25], the mechanism discussed here is qualitatively different, being driven by rare, stochastic infection events rather than by a deterministic instability. More broadly, our system relates to the physics of stochastic reaction–diffusion fronts, where demographic noise due to finite population size alters front propagation and produces fluctuations absent in deterministic descriptions [26, 27, 28, 29].

To address this gap, we develop a discrete stochastic model of phage–bacteria interactions on a two-dimensional lattice, incorporating chemotaxis and nutrient consumption. This approach tracks population counts per lattice site, rather than treating densities as continuous fields. Although individual cell trajectories are not resolved, the event-based stochastic dynamics capture essential effects of demographic noise and population discreteness that deterministic formulations miss. With this framework, we explore how parameters such as adsorption rate, burst size, and the effective local population size influence emerging spatial patterns. We show that rare, discrete infection events play a central role in generating and shaping spatial heterogeneity, a mechanism that is inaccessible to deterministic PDE models and has not been previously identified in computational models of phage–bacteria co-propagation.

## 2 Model

### 2.1 Principles of rules in the model

We consider a 2-dimensional lattice of size *L* × *L*, with each site having a linear size of Δ*x* in the xy plane. We assume the depth of the system Δ*z* is a constant and each site is locally well-mixed in the volume *V*_*site*_ = Δ*x*^2^Δ*z*. This assumption requires that bacteria and phages can diffuse across Δ*z* within a time step, which may not hold in thick agar layers. Thus, Δ*z* parametrizes the effective local population size; when we vary it to probe the role of stochasticity, large values should be understood as a theoretical limit rather than a literal agar thickness. For each lattice point (*x, y*), we consider the number of uninfected bacteria *H*(*x, y*), the number of infected bacteria *I*_*j*_(*x, y*) at the infection stage *j* (with *j* = 1, 2, …, *m*), the number of free phages *P* (*x, y*) as discrete variables (non-negative integers), the nutrient concentration *n*(*x, y*) and the chemoattractants concentrations *c*_*k*_(*x, y*) (with *k* = 1, 2, representing serine and aspartate, respectively [16]) as continuous variables. We update the system at a small discrete time step Δ*t*.

The principles of the simulation are strongly inspired by the PDE model by Ping et al. [21]. The two chemoattractant fields are introduced to reproduce the two distinct migrating fronts that bacterial populations exhibit in experimental assays [21]. What is essential for the model is the separation between a growth-limiting nutrient and one or more lower-concentration chemoattractants that serve as navigation cues, rather than treating the nutrient itself as the chemoattractant, as emphasized in [16].

Below, we summarize the basic principles of the model. The algorithm and default parameter values are summarised respectively in the Supplementary Materials S1 and S2 and a schematic representation of the model is shown in Figure 1.

**Figure 1.**
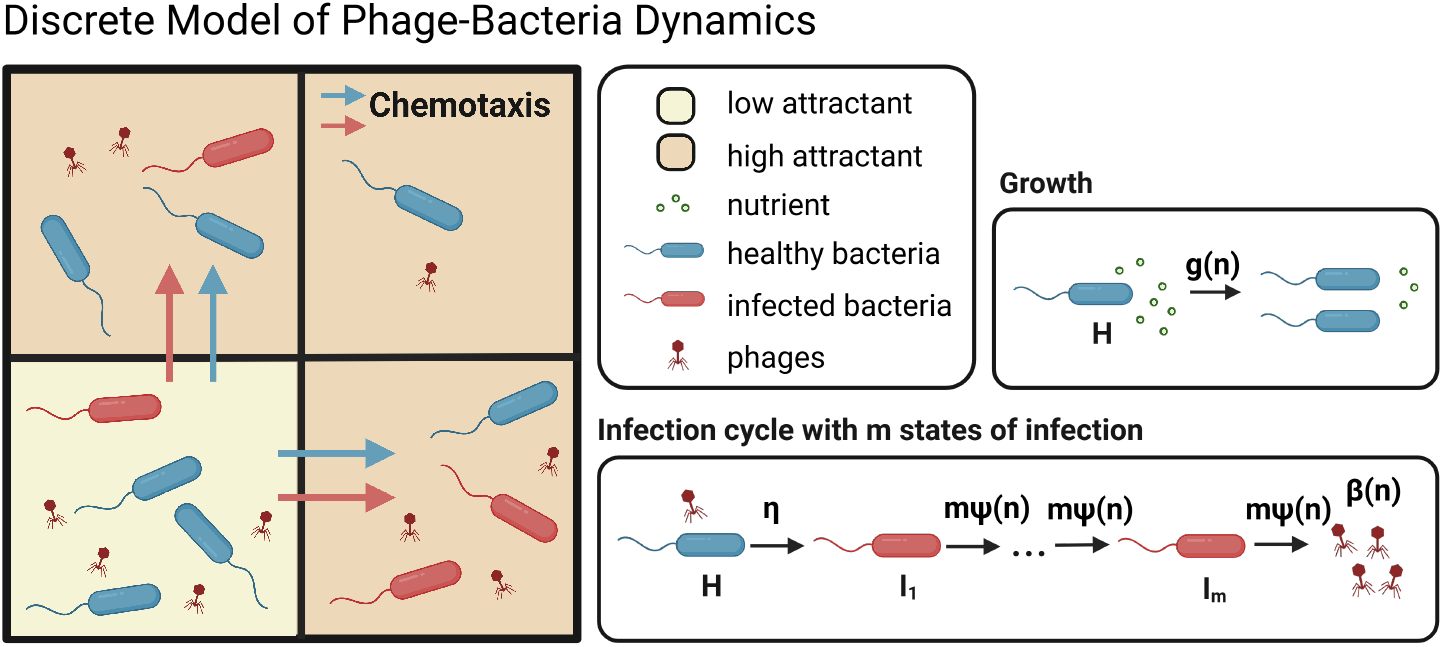
Schematic representation of the stochastic phage-bacteria model. Healthy bacteria grow by consuming nutrients and move via chemotaxis in response to attractant gradients. Phages infect bacteria, initiating a multi-step infection cycle with *m* states (*I*_1_, *I*_2_,…, *I*_*m*_) that culminates in lysis and phage release. Created in BioRender. https://BioRender.com/c7eog9f

### Bacterial Growth and Nutrient Dynamics

The dependence of the bacterial growth rate *g* on the local nutrient concentration *n*(*x, y*) is modeled using the Monod growth law [30]:

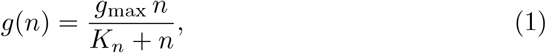

where *g*_max_ represents a constant maximum growth rate, and *K*_*n*_ is the half-saturation constant. We treat *g*(*n*) as the healthy cell’s division rate: concretely, each healthy bacterium divides with probability *g*(*n*) Δ*t* per time step, hence the number of division events happening to *H* healthy cells at a given site is drawn from a binomial distribution.

In each location, the nutrient is consumed by all bacteria *B*(*x, y*) = *H*(*x, y*) + ∑_*j*_ *I*_*j*_(*x, y*) with a consumption rate proportional to *g*(*x, y*), with the yield coefficient *Y* . Chemoattractants are assumed to also be consumed and diffuse in a similar manner. If all the field were continuous, they should obey

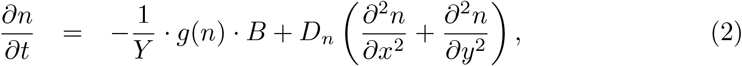

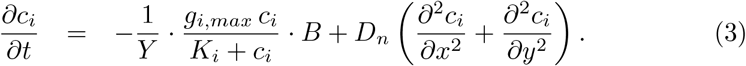

Here, *D*_*n*_ is the nutrient/chemoattractant diffusion constant, and *g*_*i,max*_ and *K*_*i*_ are chemoattractant *i*’s maximum consumption rate and Michaelis-Menten constant, respectively.

### Diffusion and Chemotaxis

Phages are assumed to diffuse at a diffusion constant *D*_*p*_, while bacteria move diffusively due to run-and-tumble motion [31] with the diffusion constant *D*_*b*_, when there is no chemoattractant gradient.

In addition, when there is gradient in chemoattractants, bacteria (both healthy and infected) bias their movements according to the gradient of a log-ratio function of the chemoattractants

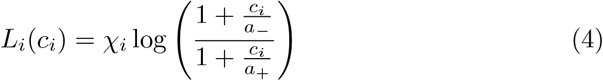

where *χ*_*i*_ is the chemotactic sensitivity for the *i*-th attractant, and *a*_−_ and *a*_+_ are the saturation constants [**tu2008modelling**, 16]. These effects are implemented as stochastic redistribution of bacteria and phages in their neighbouring sites.

### Infection and Lysis Dynamics

The infection is modeled by assuming that phages are able to adsorb to both healthy and infected bacteria, with the same adsorption rate *η*. It is implemented as a stochastic process in which individual phages adsorb to bacteria at each lattice site. During each time step, a phage adsorbs to healthy bacteria with probability proportional to the local number of susceptible hosts, *η H*(*x, y*) Δ*t*, and similarly to already infected bacteria with probability *η* ∑_*j*_ *I*_*j*_(*x, y*) Δ*t*. Adsorption events are sampled from binomial distributions, and successful adsorption to healthy bacteria initiates infection, while adsorption to infected bacteria removes phages without creating new infections. To represent the latent period between phage adsorption and host lysis, we model infected bacteria as progressing through *m* discrete infection stages, *I*_*j*_ with *j* = 1, 2,…, *m*, before bursting. The infection stage progress to the next stage at a constant rate, with each stage having an average lifetime 1*/*(*mψ*(*n*)). The nutrient-dependent lysis rate *ψ*(*n*) is modeled as

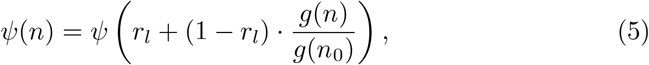

where *ψ* is the maximum lysis rate, *r*_*l*_ is the ratio between the minimum and maximum lysis rates, *n*_0_ is the initial nutrient concentration, and *g*(*n*) is the local bacterial growth rate [21].

The cells in the infection state *m* will burst to produce phage progeny at the same rate *mψ*(*n*). The burst size of the lysing bacteria also depends on the nutrient concentration:

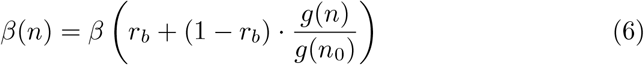

where *β* is the maximum burst size, and *r*_*b*_ is the ratio of the minimum burst size to its maximum [21].

These formulations capture the experimentally observed dependence of the latent period and the burst size on host physiological state, such that lower host growth rates extend the latent period and reduce burst size [32]. Introducing *m* successive infection stages corresponds to modelling the latent period distribution as an Erlang distribution. The total mean latent time is fixed to be 1/*ψ*(*n*), while the variability is controlled by *m*: the coefficient of variation of the Erlang distribution scales as 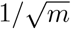. In the limiting case of *m* = 1, the latent period is exponentially distributed, corresponding to a process with maximal variance, whereas increasing *m* progressively concentrates lysis events around the mean. This formulation is useful in the present model because it allows us to isolate the role of latent-period variability in shaping emergent spatial patterns. Moreover, representing biological delays through multi-step processes is a standard technique in stochastic population and epidemic models, and provides a more realistic description than a single exponential waiting time, which tends to over-represent very short delays [10, 33]. Progresses between the states are implemented as a Poisson process happening at the given rates.

### 2.2 Initial conditions, boundary conditions, and parameters

Parameters are estimated based on the experimentally known values [21] as described in Supplementary Material S2, and the default values are summarized in the Supplementary Table S1. We simulate the system under reflective boundary conditions. At time zero, *P* (0) phages and *B*(0) bacteria are placed in the left bottom corner lattice site, and nutrients and chemoattractants *n*(0), *c*_1_(0), and *c*_2_(0) are uniformly distributed in all the lattice sites. This results in simulating 1/4 of the entire system. To verify that the reduced domain with reflective boundaries does not introduce artifacts, we repeated the simulations on the full domain and obtained consistent results (see Supplementary Figure S7).

## 3 Results

### 3.1 Emergence of spatio-temporal patterns

We simulated pattern formation from a localized inoculum of bacteria and phages using a discrete model that includes bacterial growth, nutrient and attractant diffusion and consumption, chemotactic migration, phage infection, and lysis (Figure 1; see Model section for details). Ping et al. [21] reported that the killing front advances faster than expected from free phage diffusion alone and proposed that infected bacteria remain motile and chemotactic, thereby transporting phages until lysis. We incorporate this mechanism in our model. In the absence of phage infection, the discrete model reproduces the quantitative profiles of the corresponding deteministic PDE model, with a slightly reduced leading edge due to the low-density cutoff imposed by integer population counts, consistent with the analysis by Brunet and Derrida [26, 27] (Supplementary Material S3). The model also reproduces the experimentally reported phase-diagram trends relating phage inoculum and adsorption rate (altered experimentally by changing the calcium concentration in the medium) to front radius and ring width [21] (Supplementary Material S7).

With this validation in place, we focus on a feature specific to the discrete stochastic model: killing-front roughness. A representative outcome with the default parameter set is shown in Figure 2. In this case, the number of infected states was set to *m* = 2, which produces a relatively broad distribution of latent periods with a high probability of short delays. The simulation shows the emergence of irregular killing fronts, and allow us to identify the stochastic mechanisms underlying their formation.

**Figure 2.**
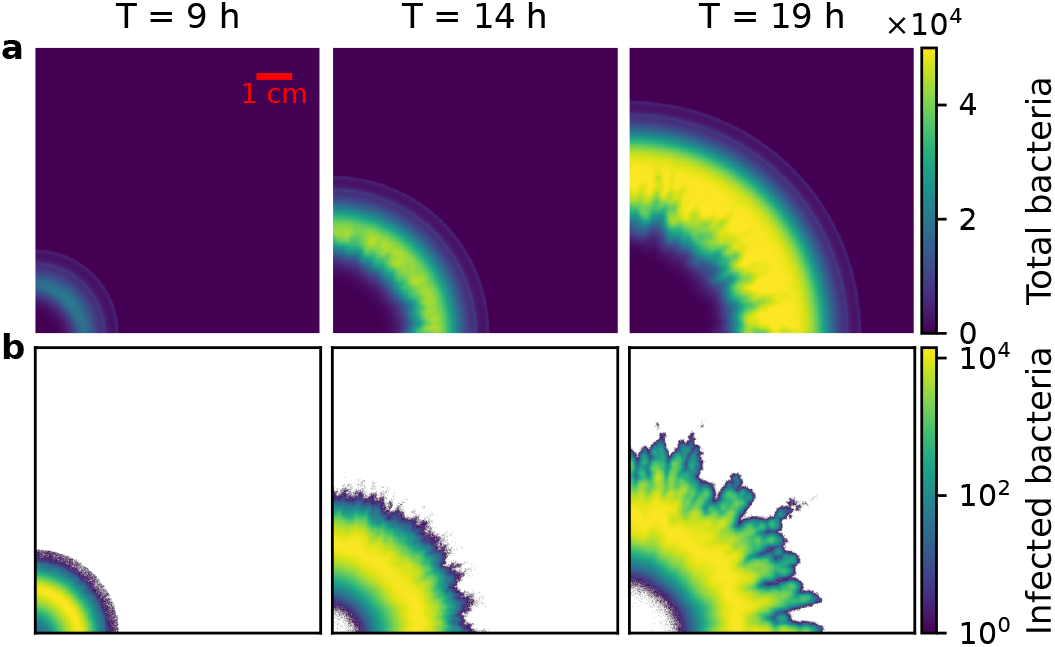
Spatio-temporal evolution of phage-bacteria dynamics. In the upper panels we show total bacteria patterns after 9 h, 14 h, and 19 h, showing the formation of irregular killing fronts. In the lower panels we show the corresponding logarithmic counts of infected bacteria, highlighting stochastic seeding events at the advancing front. Simulations start with Δ*z* × 10^4^ initial bacteria and the same amount of initial phages in the lower-left corner; depth Δ*z* = 5 mm, *β* = 20 and *η* = 1.5 · 10^−8^ mL h^−1^.

The roughening of the front arose from stochastic infection events. As shown in the bottom panel of Figure 2, where log(∑_*j*_ *I*_*j*_(*x, y*)) highlights infected sites, even a single infected bacterium at the leading edge could seed a new extension. If such a cell migrated sufficiently far before lysis, its lysis in in the middle of chemo-taxing sensitive cell population initiated a new cluster of infections that enabled phages to keep pace with the expanding bacterial front. In regions where no such rare events occurred, phages lagged behind and the infection front advanced more slowly.

The emergent patterns depended strongly on the phage–bacteria interaction parameters and initial conditions. To probe how infection dynamics shapes these patterns, and in particular when stochastic roughening emerges, we set out to systematically vary the adsorption rate (*η*) and burst size (*β*) across a broad range of values (Figure 3).

**Figure 3.**
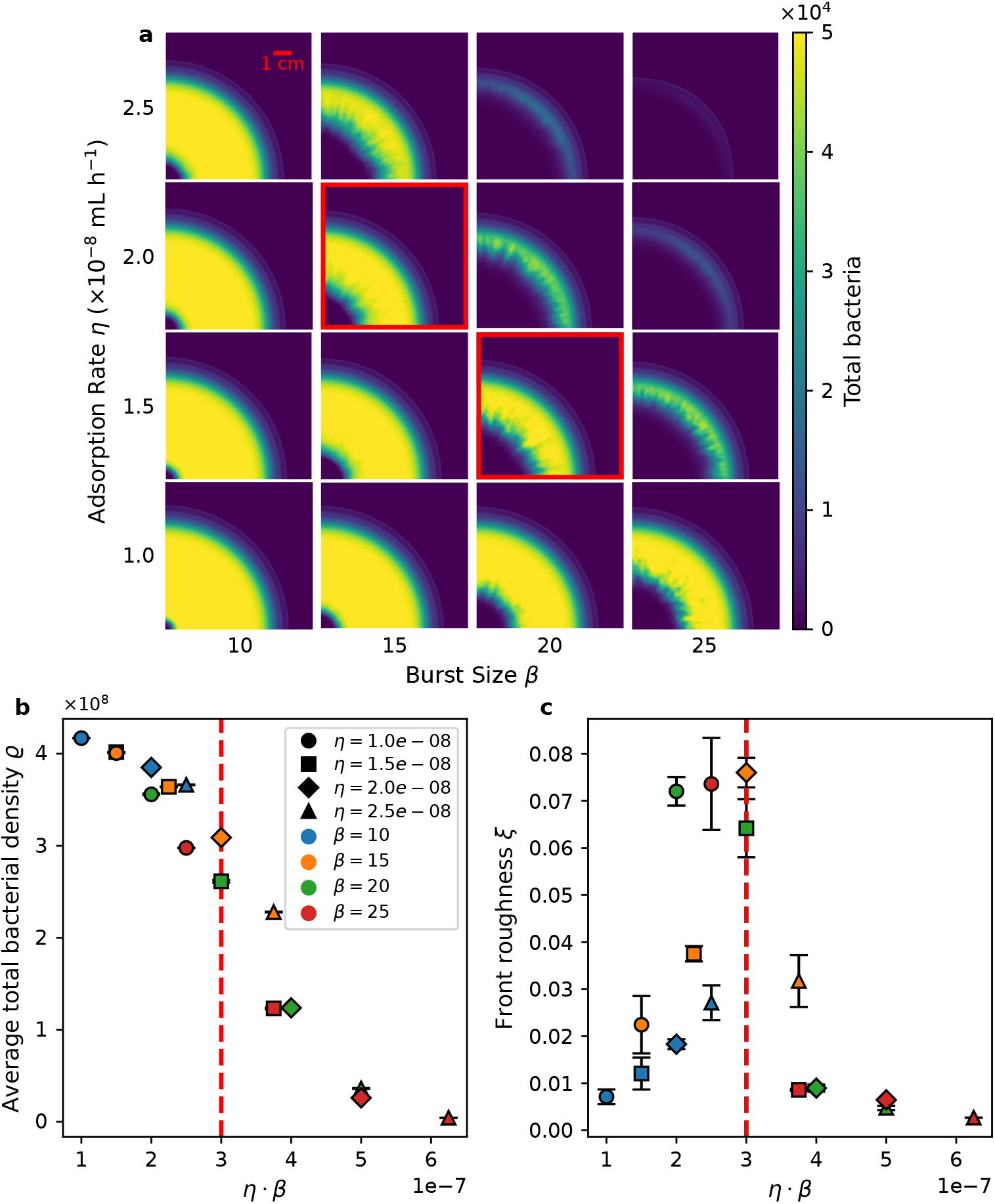
Dependence of spatial dynamics on phage–bacteria interaction parameters. (a) Phase diagram of final bacterial patterns for varying *η* and *β*. (b) Average total bacterial density *ρ* after 19 h as a function of *η* · *β*. Each symbol represents the mean across *n* = 3 independent replicates, with error bars indicating the standard error of the mean (Most of the cases error is too small to be visible). (c) Frontier roughness ratio *ξ* as a function of *η* · *β*. Each symbol represents the mean across *n* = 3 independent replicates, with error bars indicating the standard error of the mean. The dashed red lines mark the parameter combinations highlighted in panel (a). Simulations start with Δ*z* · 10^4^ initial bacteria and the same amount of initial phages in the lower-left corner; Δ*z* = 5 mm.

For small *η* and *β*, phage killing was weak: bacterial migration dominated, and phages were left behind, remaining only near the inoculation site. As *η* and *β* increased, the bacterial front became progressively suppressed, culminating in collapse of the migrating population. Stochastic, irregular patterns arose at intermediate values, where phage expansion and bacterial migration proceeded at comparable rates. In this regime, rare infection events determined whether phages successfully colonized new regions, underscoring the importance of discreteness in shaping spatial dynamics.

To assess whether deterministic dynamics alone can generate similar irregular fronts, we also simulated the corresponding two-dimensional deterministic PDE model. In contrast to the discrete stochastic simulations, the PDE model consistently produces smooth, radially symmetric infection fronts, with no sign of front instability (Supplementary Material S3, Figures S1 and S2). Since Ping *et al*. suggested the spatial variations of initial phage inoculum as a possible source of the stochasticity in the killing front [21], we also simulated a case where a strong spatial heterogeneity was introduced in the initial phage inoculum. We found that the deterministic PDE dynamics rapidly smooth out these inhomo-geneities, and the infection front remains compact and circular at later times (Supplementary Material S6). These results indicate that neither deterministic reaction–diffusion dynamics nor inoculum heterogeneity alone are sufficient to generate persistent front roughness, highlighting the essential role of stochastic infection events.

To quantify bacterial survival in the parameters sweep simulations, we calculated the average bacterial concentration at the end of each run (19 h after inoculation), defined as

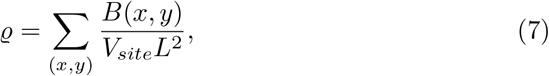

where *B*(*x, y*) is the number of bacteria at lattice site (*x, y*), *V*_site_ is the site volume, and *L*^2^ the total system area. The resulting values of *ρ* are shown in Figure 3b for each (*η, β*) pair. When plotted against the product *η* · *β*, the results collapse onto a single curve, indicating that the combined efficiency of adsorption and phage production acts as a natural control parameter. To illustrate representative cases within the intermediate regime, we selected two parameter combinations (*η, β*) corresponding to distinct realizations of stochastic patterns, highlighted in red in Figure 3a. In Figure 3b a dashed line is drawn at the associated values of *η* · *β* to indicate their position within the parameter space. Both cases exhibit irregular, stochastic patterns, characteristic of this intermediate regime.

To quantify geometric irregularity of the infection front, we define a dimensionless roughness measure:

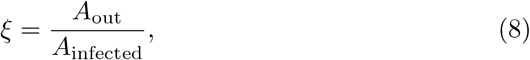

where where *A*_infected_ is the total area of the infected region and *A*_out_ is the portion of that region lying outside a reference quarter circle constructed to have the same area *A*_infected_. This measure therefore quantifies how strongly the infected domain deviates from a compact circular shape, with larger *ξ* corresponding to more irregular, protrusive fronts (see Supplementary Material S4 for details). The dependence of *ξ* on the combined parameter *η* · *β* is shown in Figure 3c. We find that *ξ* peaks at intermediate values of *η* · *β*, indicating the emergence of irregular, stochastic invasion dynamics, consistent with the parameter region highlighted in Figure 3a. In contrast, for both small and large *η* · *β, ξ* approaches zero, corresponding to compact, approximately circular infection fronts.

### 3.2 The irregular pattern is due to the small number effect

To quantify the role of stochasticity—particularly rare events in which single infected bacteria migrate far ahead of the front—we varied the model depth parameter Δ*z*. In the model, Δ*z* does not represent the physical agar thickness; rather, it defines an effective vertical extent used to convert concentrations into discrete particle numbers at each lattice site. We assume the system is well mixed along this depth direction irrespective of its value, so the dynamics remain effectively two-dimensional. If the physical agar thickness were changed, the well-mixed assumption would break upon certain thickness where mixing by diffusion over one time step become insufficient. Initial conditions were rescaled with Δ*z* to keep all concentrations fixed. Increasing Δ*z* therefore increases the number of individuals per lattice site without changing concentrations or transport parameters, thereby reducing demographic noise. In the limit Δ*z* → ∞, fluctuations arising from finite individual numbers vanish by the law of large numbers.

Changing the effective number of individuals alters both the likelihood and the impact of rare events. Larger depths increase the chance that a single infected cell travels far for each location, but the effect of such an event becomes diluted because each burst affects a smaller fraction of the local population. This trade-off is evident in Figure 4. As shown in Figure 4a, greater depths produce more radial “spikes” in the phage front, but the spikes appear with weaker contrast, yielding smoother phage fronts and more uniform bacterial distributions overall. The dependence of the front roughness *ξ* on Δ*z* is shown in Figure 4b: systems with lower depth exhibit higher *ξ* values, indicating rougher, more irregular fronts, while increasing Δ*z* leads to smoother, more circular colonies, characterized by lower values of *ξ*.

**Figure 4:**
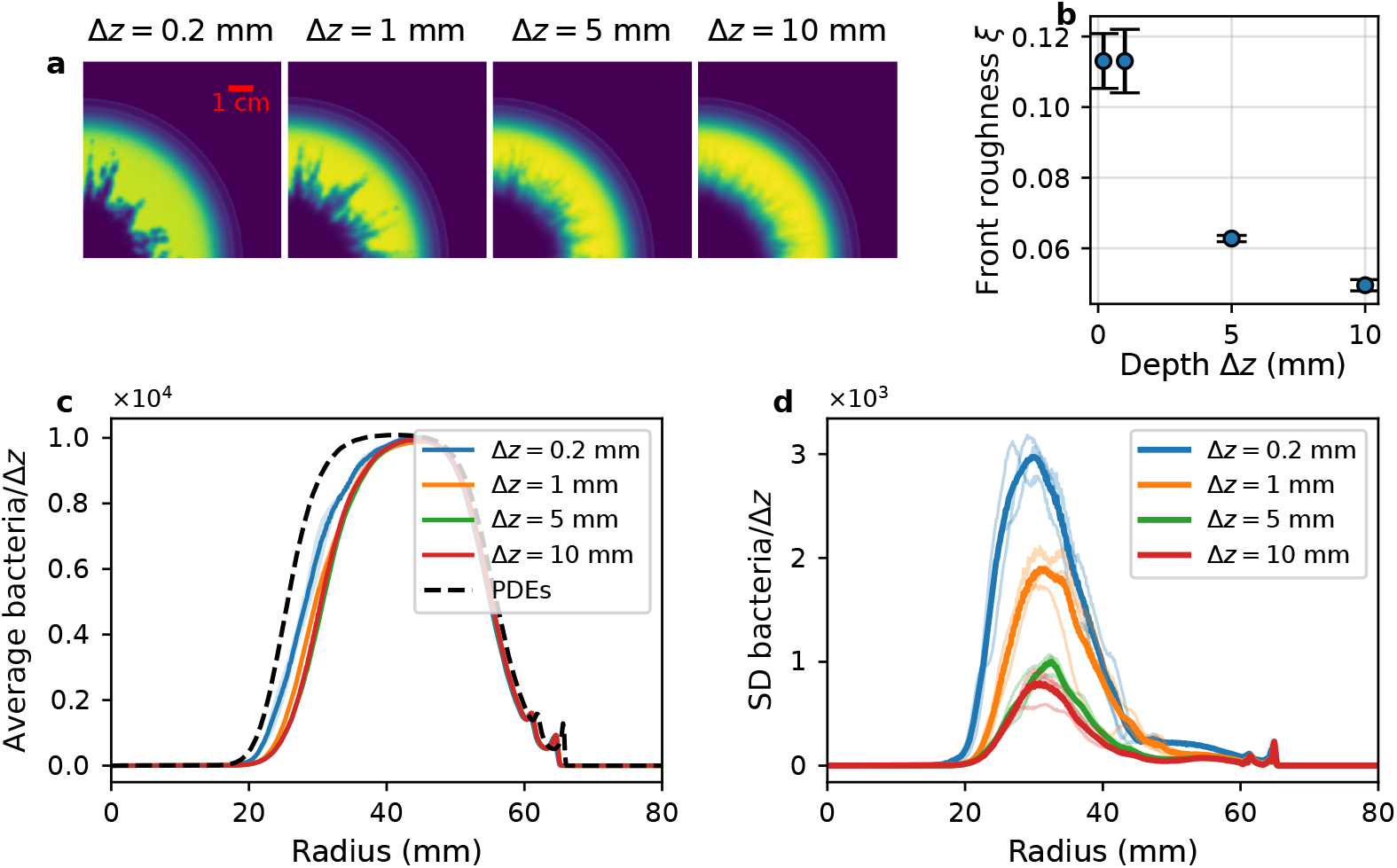
Impact of system depth on phage–bacteria infection dynamics. (a) Final patterns at 19 hours for simulations with different depths: Δ*z* · 10^4^ initial bacteria and the same amount of initial phages in site (0, 0). Simulations use *β* = 20 and *η* = 1.5 · 10^−8^ mL h^−1^. (b) Frontier roughness ratio *ξ* as a function of Δ*z*. Each point shows the mean ± standard error of the mean across *n* = 3 independent replicates. (c) Radial average profiles (radius in mm) for each depth (Δ*z*). Curves show the mean across n=3 independent replicates, with shaded bands indicating the standard error of the mean across replicates. To enable comparison across depths, the mean is normalized by Δ*z* (y-axis: Average bacteria/Δ*z*). A dashed black curve shows the corresponding PDE result for reference. (d) Radial standard-deviation profiles. Thin lines indicate individual replicate standard deviation profiles, and bold lines show the pooled standard deviation (square root of the mean of replicate variances). Values are normalized by Δ*z* (y-axis: Std bacteria/Δ*z*).

This trend is further quantified in Figure 4c, which shows radial averages, and in Figure 4d which shows standard deviations of bacterial densities. At larger depths, the radial profiles become smoother and the standard deviations decrease, indicating reduced spatial heterogeneity. For reference, we compare these results to the corresponding deterministic PDE model, which produces a smooth and stable infection front (Supplementary Material S3). However, the ensemble average at large depth does not lie closer to the PDE curve for larger Δ*z* within the range we could simulate: at small depths, rare runner events are infrequent, so the mean is often dominated by typical ‘no-runner’ realizations and can appear closer to the PDE. At large depths, runner events become common but individually weaker, which lowers variability yet can bias the mean killing front outward. We could not simulate much larger Δ*z* due to the high computational cost, hence we could not confirm or disconfirm if the system eventually converge to PDE result in Δ*z* → ∞ limit.

### 3.3 Effects of latent period length distribution on infection front stochasticity

The results so far indicate that irregular front roughening originates from the amplification of rare events where a single infected bacterium lyses ahead of the average front. For such an event to produce a visible clearing spike, the initial burst must be amplified by successive rounds of infection before the major killing front catches up. This process may couple two extremes of the latent period distribution: a bacterium with a relatively long latent period can carry phages far ahead before bursting, while progeny from a bacterium with a short latent period can rapidly initiate secondary infections. Consequently, the distribution of latent periods is expected to strongly influence the emergent patterns.

To test this, we varied the number of discrete infected states *m*, which generates an Erlang distribution of latent period length with coefficient of variation 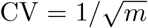. When *m* = 1, lysis times follow an exponential distribution (CV = 1), leading to highly irregular fronts with broad, dark spikes (Figure 5, leftmost panel). As *m* increases, the latent period distribution becomes more sharply peaked around the mean. This reduction in variance suppresses extreme outliers and diminishes the large-scale roughening of the front. Panels for *m* = 2 and *m* = 3 show progressively thinner, less pronounced spikes, and by *m* = 5 the infection front appears smooth and uniform.

**Figure 5.**
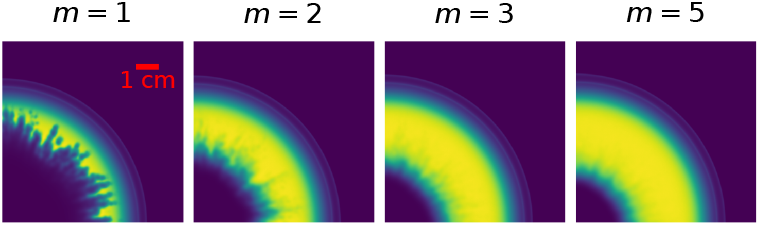
Effect of lysis time distribution on spatial infection patterns. Infection patterns after 19 h for Erlang distributions with *m* = 1, 2, 3, 5 substeps (coefficient of variation 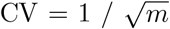). As *m* increases, the phage spread decreases and the infection front becomes smoother. All simulations use *β* = 20 and *η* = 1.5 · 10^−8^ mL h^−1^.

We further tested whether adsorption kinetics could compensate for this effect. As shown in Figure S11, increasing *η* at *m* = 5 did not restore the thick, irregular spikes characteristic of the low-*m* regime. Taken together, these results indicate that the shape of the latent period distribution—not just the mean latent time or adsorption rate—critically determines macroscopic pattern variability. Exponential lysis amplifies demographic noise and produces rough, irregular fronts, whereas multi-step Erlang distributions concentrate lysis events near the mean, suppressing outliers and yielding smoother, more deterministic fronts.

As we stated before, increasing the parameter *m* effectively narrows the distribution of latent periods. However, it remains unclear whether the shorter or longer latent periods are primarily responsible for the emergence of irregular pattern. To address this, we constructed a modified model (presented in more detail in Supplementary Material S8) in which, upon infection, each cell is stochastically assigned to one of several latency-time distributions. In 90% of cases, cells follow the Erlang distribution with *m* = 3 and a mean latency of 1 hour, while the remaining 10% are assigned to a Erlang distribution with *m* = 3 and a mean latency of either 3 hours or 20 minutes, depending on the simulation. The obtained distributions are shown in Figure 6a with the Erlang distribution of *m* = 3. This approach allows us to disentangle the respective contributions of short and long latent periods to the overall population dynamics. The results, presented in Figure 6b, show that rare early lysis events are more effective at restoring roughness than late outliers, highlighting the asymmetric role of the distribution tails in shaping front geometry. A similar asymmetry arises when considering heterogeneity in motility (Supplementary Material S8): less responsive cells have little effect, whereas a minority of more motile infected cells can travel further up the gradient and lyse ahead of the front, again generating forward outliers that restore roughness.

**Figure 6.**
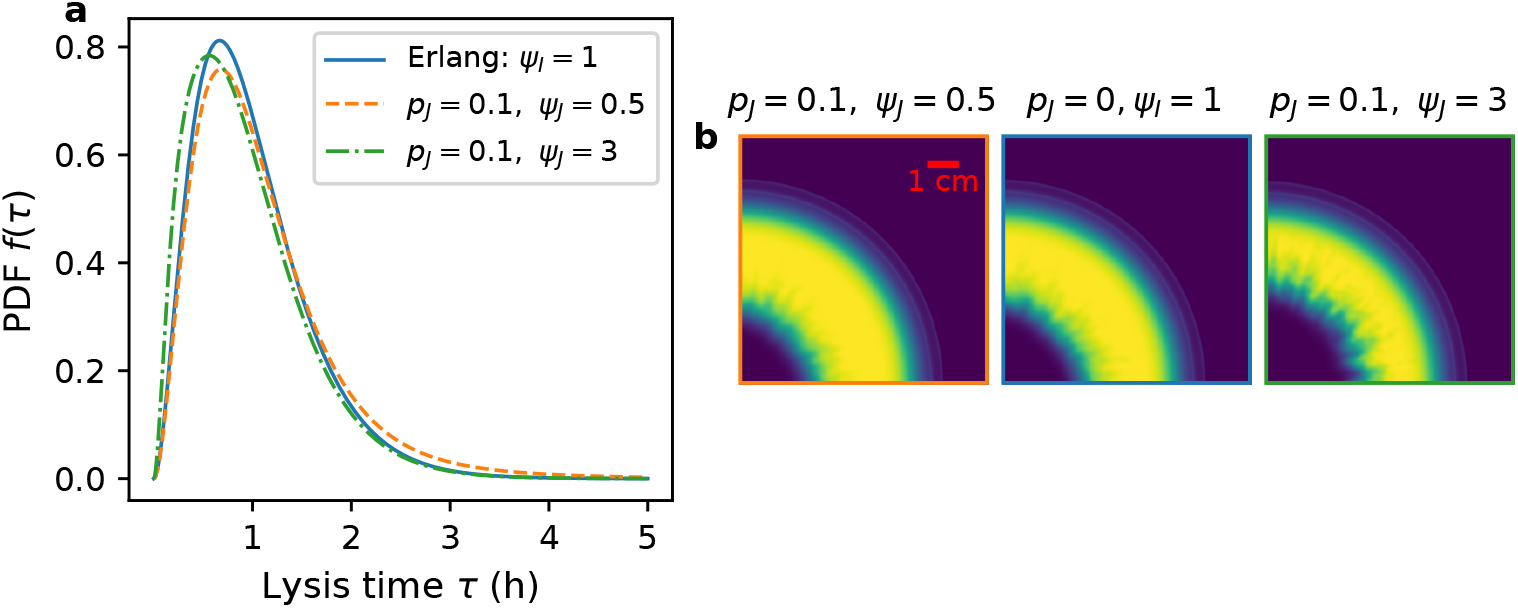
Impact of faster or slower lysing subpopulations on infection front structure. (a) Probability density functions of lysis time for an Erlang distribution (*ψ* = 1, blue) and mixtures with a minority subpopulation, having either slower lysis *ψ*_*J*_ = 0.5, orange dashed) or faster lysis (*ψ*_*J*_ = 3, green dash-dotted). (b) Spatial infection fronts under mixed lysis rates: snapshots of total bacterial density for three parameter sets: (left) minority fast-lysing infections (*p*_*J*_ = 0.1, *ψ*_*J*_ = 3, *ψ*_*I*_ = 1); (center) baseline Erlang distribution (*p*_*J*_ = 0, *ψ*_*I*_ = 1); (right) minority slow-lysing infections (*p*_*J*_ = 0.1, *ψ*_*J*_ = 0.5, *ψ*_*I*_ = 1). Rough fronts are restored when early lysis events are present (left), while late outliers (right) do not have a strong effect on the shape of the front.

## 4 Discussion

We developed a discrete stochastic model that captures qualitative features of the irregular spatial patterns observed in experimental studies of interactions between phages and motile bacteria, and reveals mechanisms that are inaccessible to deterministic PDE approaches. Rather than aiming to reproduce experimental realizations in quantitative detail, the model was designed to isolate a mechanism responsible for irregular fronts and to reveal how stochasticity at the level of individual infection events is amplified into macroscopic spatial patterns. By explicitly modelling discrete events in phage-bacteria interactions and incorporating stochasticity, the model revealed how spatial heterogeneity naturally emerges from the interplay between bacterial migration and phage infection.

In particular, we showed that rare infection events, such as a single infected bacterium migrating ahead of the front before lysis, can generate new centers of infection, having a strong effect on large-scale dynamics. This sensitivity to individual events underscores the importance of treating infection and transport as discrete processes. To our knowledge, this rare-event-driven roughening mechanism, where a single stochastic infection event at the front is amplified into a macroscopic spatial protrusion through successive rounds of infection, has not been described in previous models of phage–bacteria spatial dynamics, which have focused on deterministic PDE frameworks that inherently smooth over such events.

Our results identify the product of adsorption rate and burst size (*η* · *β*) as a key control parameter governing the killing front morphology in the model. Since *η* governs how quickly phages encounter hosts and *β* characterizes the number of progeny per lysis, their product sets the per-cycle amplification of the infection cascade and thus the speed at which the killing front can advance. As *η* · *β* increases, the simulations shift from expanding bacterial fronts to phage-driven collapse, reflecting a trade-off between bacterial dispersal and phage reproductive efficiency. This behavior is qualitatively consistent with the experiments of Ping et al. [21], where tuning the P1_vir_ adsorption rate via calcium concentration leads to corresponding changes in spatial patterning, with irregular fronts observed in an intermediate regime between bacterial- and phage-dominated dynamics. A quantitative comparison of front roughness with the experiments is not feasible from the published figures alone, which provide only a single detailed rough-front example and insufficient geometric resolution for applying our roughness metric. We therefore restrict the comparison to qualitative features. We demonstrate that increasing the depth of the lattice, which effectively increases the number of individuals per site, suppresses the impact of stochastic fluctuations and leads the system toward a smooth killing pattern. This suppression of roughness with increasing population size is analogous to the finite number corrections to stochastic front propagation [26, 27], where discreteness effects vanish as the effective population size grows. However, our system differs from the single-species case in that the noise acts on a secondary infection front propagating within a primary chemotactic wave. Overall, our result suggests that population size and demographic noise fundamentally shape infection front dynamics, with implications for understanding pattern formation in natural microbial communities.

Finally, by varying the number of discrete infected states, we show that the shape of the latent period distribution itself—not just its mean—critically modulates front roughness. We demonstrated that the occasional burst with short latent time is particularly important for the irregular killing patterns, likely due to exponential amplification of phage following after such an event. The asymmetric role of the distribution tails, with rare early-lysis events having a disproportionate effect on front roughness compared to late-lysis outliers, is consistent with the established principle that stochastic front dynamics are dominated by rare fluctuations at the leading edge [26, 29]. This is an experimentally testable prediction if one can identify phage mutants that have different latent period distribution. The impact of latent period distribution has been recently highlighted in several theoretical works [34, 35]. This indicates the importance of characterizing the latent time distributions experimentally, which is becoming feasible thanks to the recent development of single cell measurements [36].

We have also found that having subpopulation of infected cell that swim faster than others could also be a possible way to have irregular pattern. It should be noted that the infection itself is more likely to reduce bacterial motility by interfering with cellular physiology; however, a similar effect is expected to arise if the bacterial population already exhibits heterogeneity in chemotactic ability prior to infection. It should also be noted that, even though some degree of swimming of infected bacteria is widely accepted based on the experimentally observed pattern [21, 22, 24], a recent single-cell experiment showed infection by *λ* phage can diminish the ability of *E. coli* flagella motor to rotate [37]. Furthermore, a previous study has reported that phages can be transported by non-host motile bacteria [38], suggesting that transport by uninfected cells may also play an important role. Characterizing attachment of phage to swimming bacteria and the ability of infected cells to swim at single-cell level would also be an important experimental step to fully understand the observed killing patterns.

The model has several limitations. Populations are tracked as integer counts per lattice site rather than as individual cells, so processes that depend on cell-level trajectories or infection histories are not resolved. Motility and chemotactic parameters are assumed identical for all bacteria; in reality, phenotypic variability can produce subpopulations with distinct swimming speeds or chemotactic responses [39, 40, 41], potentially altering infection dynamics and spatial pattern formation. Additionally, all bacteria are assumed to be susceptible to infection, and no evolution is permitted during the simulation. In natural systems, spontaneous mutations can lead to resistance or altered infection phenotypes, which are crucial for long-term coexistence and evolution [42, 43]. One could incorporate heterogeneous motility or phage resistance into the model by dividing the populations into different subcategories that has distinct chemotactic parameters or a resistance state. Such a model would allow investigation of how diversity and evolution reshape the spatial patterns described here.

Our model connects single-cell stochasticity with population-scale organization, linking to the broader framework of stochastic reaction–diffusion fronts [28] and noise-driven pattern formation in expanding microbial populations [44]. This insight may extend beyond phage–bacteria dynamics to other systems in which growth and dispersal are coupled, such as microbial range expansions [45, 46, 47], ecological invasions [48], or viral epidemics. More generally, our findings emphasize the need to consider demographic fluctuations not as noise to be averaged out, but as an intrinsic driver of spatial pattern formation in living systems.

## Code Availability

The source code will be made available at:https://github.com/laurabergamaschi22/phage_motile_bacteria_model_code.git

## Acknowledgement

This research was funded by the Novo Nordisk Foundation (NNF21OC0068775).

