## Supplemental Text for "Stochastic Emergence of Irregular Infection Fronts in Motile Bacteria-Phage Populations"

### Supplementary Materials for:

### S1 Discrete Model Simulation Algorithm

We update the system's state with discrete time step $\Delta t$. The update per time step is executed in the following order.

**1. infection.** For each lattice point $(x, y)$, the number of phages that get adsorbed by healthy bacteria, infected bacteria or that do not get adsorbed at the time step are extracted from the multinomial distribution, with the following probabilities:

$$P_h = \eta \, H(x, y) \, \Delta t, \tag{9}$$

$$P_i = \eta \sum_j I_j(x, y) \, \Delta t, \tag{10}$$

$$P_{\text{none}} = 1 - P_h - P_i, \tag{11}$$

where $P_h$ is the probability for phages to adsorb to healthy bacteria, $P_i$ is the probability to adsorb to infected bacteria and $P_{\text{none}}$ the probability to not adsorb. The phages that adsorb to healthy or infected bacteria are removed from the site, and the number of healthy bacteria that become infected is determined as the minimum between the number of healthy bacteria available at the site and the number of phages adsorbing to healthy bacteria. This approach ensures that infections do not exceed the number of susceptible hosts and is based on the assumption that each phage, on average, targets a different healthy bacterium.

**2. lysis.** The number of infected bacteria at lattice point $(x, y)$ that pass from infection step $k$ to $k + 1$, $\Delta I_k(x, y)$ is extracted from the binomial distribution, with probability of each infected cell's transition per time step being $m\psi(n(x, y)) \cdot \Delta t$. The number of infected bacteria that lyse from state $m$, $\Delta I_m(x, y)$, is also extracted from the binomial distribution, using the same probability. For each lattice site, $\Delta I_k(x, y)$ is first calculated for all $k$, and then $I_k(x, y)$ is reduced by $\Delta I_k(x, y)$. $I_{k+1}(x, y)$ is increased by $\Delta I_k(x, y)$ for $k < m$, and the number of free phage $P(x, y)$ is increased by $\beta(n(x, y))\Delta I_m(x, y)$.

**3. bacteria chemotaxis first attractant (serine).** First, we calculate the log-ratio of the first chemoattractant based on (4) at each location $(x, y)$ as $L_1(c_1(x, y))$. Then, we compute the local gradient $v_{d,1}(x, y)$ based on the value of each of the 4 nearest neighbours $d \in \{(x+1, y), (x-1, y), (x, y+1), (x, y-1)\}$ as

$$v_{d,1}(x, y) = \max\{L_1(x_d, y_d) - L_1(x, y), 0\} \tag{12}$$

where $(x_d, y_d)$ are the coordinates of the neighbour. For each population — healthy bacteria $H(x, y)$ and each infected bacteria sub-population $I_j(x, y)$ $(j = 1, \cdots, m)$ — we independently compute how many bacteria will move out of the site by sampling from the binomial distribution with probability $v_{tot}(x, y) \cdot \Delta t$, where $v_{tot,1} = \sum_d v_{d,1}(x, y)$. The number of bacteria moving in each direction is then extracted from the multinomial distribution, using probabilities $v_d/v_{tot}$. In this way, higher gradients (larger $v_d$) lead to larger fluxes of bacteria along that direction, while directions with negligible gradients receive fewer (or no) bacteria moving in that timestep. If the gradient is negative, no bacteria are moving. The movement is executed after the number of bacteria to move is calculated for all the sites.

**4. bacteria chemotaxis second attractant (aspartate)** The same algorithm as step 3 is executed, with $c_1$, $L_1$, $v_{d,1}$, and $v_{tot,1}$ replaced by $c_2$, $L_2$, $v_{d,2}$, and $v_{tot,2}$, respectively.

**5. bacteria diffusion** To model the diffusion of bacteria, the number of bacteria that move out of each site is extracted from the binomial distribution, with the probability to move to one of the 4 nearest neighbor sites $v_b \cdot \Delta t$. Here, the exit rate $v_b$ is determined by the bacterial diffusion coefficient as $v_b = 4D_b/(\Delta x)^2$. Bacteria have then equal probability to move in each of the 4 possible directions. The movement is executed after the number of bacteria to move is calculated for all the sites.

**6. phage diffusion** Similarly to bacteria diffusion, the number of phages that move out of each site is extracted from the binomial distribution, with the probability to move to one of the 4 nearest neighbor sites $v_p \cdot \Delta t$,. Here, the exit rate $v_p$ is determined by the phage diffusion coefficient as $v_p = 4D_p/(\Delta x)^2$. The movement is executed after the number of phages to move is calculated for all the sites.

**7. nutrient diffusion** The nutrients are updated according to the finite-difference methods, i.e., $n(x, y)$ changes by $D_n[(n(x-1, y) + n(x+1, y) - 2n(x, y)]/\Delta x^2 + [n(x, y-1) + n(x, y+1) - 2n(x, y))/\Delta x^2]\Delta t$.

**8. serine diffusion** Similarly to the nutrient diffusion, $c_1(x, y)$ changes by $D_n[(c_1(x-1, y) + c_1(x+1, y) - 2c_1(x, y)]/\Delta x^2 + [c_1(x, y-1) + c_1(x, y+1) - 2c_1(x, y))/\Delta x^2]\Delta t$.

**9. aspartate diffusion** The same algorithm as step 8 is executed, with $c_1$ replaced by $c_2$.

**10. nutrient consumption and growth** The nutrient $n(x, y)$ and the chemoattractants $c_i(x, y)$ are reduced by $g(n)B(x, y)\Delta t/Y$ and $[g_1 c_i/(K_i + c_i)]B(x, y)\Delta t/Y$, respectively. The number of healthy bacteria at lattice point $(x, y)$ that divide is extracted from the binomial distribution, with the probability of each healthy cell to divide being $g(n(x, y)\Delta t)$. The number of healthy bacteria $H(x, y)$ is increased according to the extracted number.

### S2 Parameters Conversion

Table S1 lists the parameters used in this study: those for the partial differential equation (PDE) model, characteristic of *E.coli* and bacteriophage $P1_{\mathrm{vir}}$ (from [1]), and those for the discrete model (DM), which represent the same system. To convert between these two parameter sets, the following quantities are useful:

- The Area of each lattice site $A = 0.01$ mm$^2$
- the Depth of each lattice site, usually $\Delta z = 5$ mm
- The Volume of each lattice site $V_{site} = \Delta z \cdot A$, converted in mL.
- The Yield Coefficient $Y = 10^8\ 1/(\mathrm{mM} \cdot \mathrm{mL})$

Some parameters remain the same for the 2 models, such as the time $T_{max}$, the growth rates $g_{max}$, $g_{1,max}$ and $g_{2,max}$, the burst size $\beta$, the latent period $\tau$ (noting that in the DM we actually use the lysis rate $\psi = 1/\tau$), and the ratios $r_b$ and $r_l$.

In the DM, we set $\Delta x = 1$ site, which means the spatial unit becomes 1 lattice site, so $A = 1$ sites$^2$. The yield coefficient is also set to 1. Bacteria and phages are represented as discrete individuals rather than concentrations, while nutrients and chemoattractants are measured in yield units. One unit of nutrient is defined as the amount required to produce one bacterium.
To convert concentrations from the PDE model (expressed in mM) to DM units, quantities are multiplied by the yield coefficient and the lattice site volume. For example:

- $n(0)[DM] = 10\ \mathrm{mM} \cdot 10^8\ 1/(\mathrm{mM} \cdot \mathrm{mL}) \cdot 5 \cdot 10^{-5}\ \mathrm{mL} = 50000$

The same process is done for $c_1(0), c_2(0), K_n, K_1, K_2, a_-, a_+$.
To convert from the PDE model's spatial units (mm) to the DM's lattice sites, diffusion and chemotactic coefficients must be adjusted by dividing by the lattice site area. For example:

- $D_b[DM] = \frac{0.05\ \mathrm{mm}^2/\mathrm{h}}{0.01\ \mathrm{mm}^2} = 5\ \mathrm{sites}^2/\mathrm{h}$

For bacterial and phage diffusion in the DM we use exit rates, defined as $v = 4 \cdot D/\Delta x^2$.
The initial concentrations of bacteria and phages are converted into initial discrete numbers by multiplying by the lattice site volume. For example:

- $P(0)[DM] = 10^9\ \mathrm{mL}^{-1} \cdot 5 \cdot 10^{-5}\ \mathrm{mL} = 5 \cdot 10^4$

The adsorption rate is converted by dividing by the lattice site volume:

- $\eta[DM] = \frac{3 \cdot 10^{-8}\ \mathrm{mL\ h}^{-1}}{5 \cdot 10^{-5}\ \mathrm{mL}} = 0.0015\ \mathrm{h}^{-1}$

Table S1: Model Parameters for PDE and DM

| **Parameter** | **Description** | **PDE** | **DM** |
|---|---|---|---|
| $L$ | Domain size | 80 mm | 800 sites |
| $\Delta x$ | Site size | 0.1 mm | 0.1 mm = 1 site |
| $\Delta z$ | Depth of site | 5 mm | 5 mm |
| $\Delta t$ | Time-step | adaptive | $2{\cdot}10^{-4}$ h |
| $T_{max}$ | Total simulation time | 19 h | 19 h |
| $n(0)$ | Initial nutrient | 10.0 mM | 50000 |
| $c_1(0)$ | Initial serine | 0.2 mM | 1000 |
| $c_2(0)$ | Initial aspartate | 0.2 mM | 1000 |
| $g_{max}$ | Max. growth rate | 2.8 $h^{-1}$ | 2.8 $h^{-1}$ |
| $g_{1,max}$ | Max. depletion rate for serine | 2.8 $h^{-1}$ | 2.8 $h^{-1}$ |
| $g_{2,max}$ | Max. depletion rate for aspartate | 0.28 $h^{-1}$ | 0.28 $h^{-1}$ |
| $Y$ | Yield coefficient | $10^8$ 1/(mM·mL) | 1 |
| $K_n$ | Monod constant for nutrient | 10 mM | 50000 |
| $K_1$ | Monod constant for serine | 0.05 mM | 250 |
| $K_2$ | Monod constant for aspartate | 0.005 mM | 25 |
| $\chi_1$ | Chemotactic coeff. for serine | 1.5 $mm^2$ $h^{-1}$ | 150 $h^{-1}$ |
| $\chi_2$ | Chemotactic coeff. for aspartate | 0.8 $mm^2$ $h^{-1}$ | 80 $h^{-1}$ |
| $a_-$ | Low chemoattractant sensing constant | 0.0035 mM | 17.5 |
| $a_+$ | High chemoattractant sensing constant. | 1 mM | 5000 |
| $D_b$ | Bacterial diffusion coeff. | 0.05 $mm^2$ $h^{-1}$ | 5 $sites^2$ $h^{-1}$ |
| $D_p$ | Phage diffusion coeff. | 0.04 $mm^2$ $h^{-1}$ | 4 $sites^2$ $h^{-1}$ |
| $D_n$ | Nutrient diffusion coeff. | 3 $mm^2$ $h^{-1}$ | 300 $sites^2$ $h^{-1}$ |
| $\eta$ | Adsorption rate | 1.0-$2.5{\cdot}10^{-8}$ mL $h^{-1}$ | 0.0002-0.0005 $h^{-1}$ |
| $\beta$ | Burst size | 10-25 | 10-25 |
| $\tau$ | Latent period | 1 h | 1 h |
| $r_b$ | Ratio of min/max burst size | 0.1 | 0.1 |
| $r_l$ | Ratio of min/max latent period | 0.5 | 0.5 |
| $m$ | Number of infected states | 2 | 2 |
| $P(0)$ | Initial phage density | $10^9$ $mL^{-1}$ | $5{\cdot}10^4$ |
| $B(0)$ | Initial bacterial density | $10^9$ $mL^{-1}$ | $5{\cdot}10^4$ |

### S3 Deterministic Partial Differential Equations Model

To complement the stochastic, discrete simulations, we implemented a deterministic continuous model in which susceptible bacteria $H$, $m$ successive infected states $I_1, \ldots, I_m$, free phage $P$, nutrient $n$, and attractant fields $c_1, c_2$ evolve by coupled reaction–diffusion equations. Infection latent period is represented by $m$ states of mean duration $\tau/m$, yielding an Erlang distribution of total latent period length with coefficient of variation $1/\sqrt{m}$. The equations are:

$$
\begin{aligned}
\frac{\partial H}{\partial t} &= D_B \nabla^2 H - \nabla \cdot \left[ H\,\chi_1 \nabla \ln \frac{1 + c_1/a_-}{1 + c_1/a_+} + H\,\chi_2 \nabla \ln \frac{1 + c_2/a_-}{1 + c_2/a_+} \right] \\
&\quad + g_{\max} \frac{n}{n + K_n} H \;-\; \eta\, H\, P, \\
\frac{\partial I_1}{\partial t} &= D_B \nabla^2 I_1 - \nabla \cdot \left[ I_1\,\chi_1 \nabla \ln \frac{1 + c_1/a_-}{1 + c_1/a_+} + I_1\,\chi_2 \nabla \ln \frac{1 + c_2/a_-}{1 + c_2/a_+} \right] \\
&\quad + \eta\, H\, P \;-\; \frac{m}{\tau} I_1, \\
\frac{\partial I_i}{\partial t} &= D_B \nabla^2 I_i - \nabla \cdot \left[ I_i\,\chi_1 \nabla \ln \frac{1 + c_1/a_-}{1 + c_1/a_+} + I_i\,\chi_2 \nabla \ln \frac{1 + c_2/a_-}{1 + c_2/a_+} \right] \\
&\quad + \frac{m}{\tau} \big( I_{i-1} - I_i \big), \quad i = 2, \ldots, m, \\
\frac{\partial P}{\partial t} &= D_P \nabla^2 P + \beta \frac{m}{\tau} I_m - \eta\, P \Big( H + \sum_{i=1}^{m} I_i \Big), \\
\frac{\partial n}{\partial t} &= D_n \nabla^2 n - \frac{g_{\max}}{Y} \frac{n}{n + K_n} \Big( H + \sum_{i=1}^{m} I_i \Big), \\
\frac{\partial c_j}{\partial t} &= D_n \nabla^2 c_j - \frac{g_{j,\max}}{Y} \frac{c_j}{c_j + K_{m,j}} \Big( H + \sum_{i=1}^{m} I_i \Big), \quad j = 1, 2,
\end{aligned}
\tag{13}
$$

These equations were discretized on a two-dimensional lattice with no-flux boundaries, using an explicit finite-difference stencil and an adaptive time step, following the approach of [2].

When evolved to $t = 19$ h, the PDE model produces a smooth, circular infection front, shown in Figure S1, without the roughening observed in the stochastic simulations, confirming that front roughness in the discrete model arises from rare events rather than from deterministic kinetics or movement.

In Figure S2 we show the infection patterns in the $(\eta, \beta)$ parameters space for the PDE model, to compare to Figure 3 of the main text. In the stochastic model rare events act like scouts that push the front forward locally, creating new outbursts of infection. Without that mechanism, the deterministic wave advances at its mean speed, never overshooting into new territory. Thus, comparing the two models at fixed time, the deterministic PDE will always show a smaller infected region than the corresponding stochastic simulation with the same parameter values.

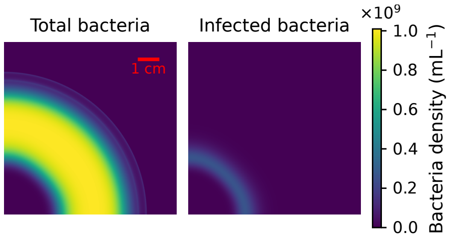


Figure S1: Deterministic PDE simulation showing smooth circular infection fronts. Deterministic PDE results at $t = 19$ h, with $\beta = 20$ and $\eta = 1.5 \cdot 10^{-8}$ mL h$^{-1}$ Left: total bacterial density $B + \sum_i I_i$, showing a circular invasion front. Right: total infected density $\sum_i I_i$, confined to a narrow ring with no stochastic branches.

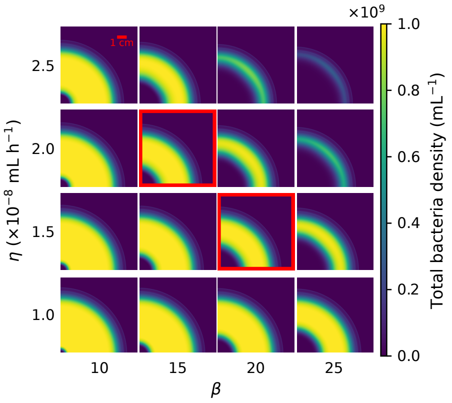


Figure S2: Phase diagram in the $(\eta, \beta)$ plane for the deterministic PDE model. The deterministic PDE simulations show no stochastic effects within the explored parameter space, in contrast with the discrete simulations.

To validate that the discrete stochastic model reproduces the PDE model in absence of stochastic infection dynamics, we performed simulations in which phage infection and lysis were disabled ($\eta = 0$), leaving only bacterial diffusion, chemotaxis, growth, and nutrient consumption. Figure S3 compares radially averaged bacterial density profiles obtained from the discrete model and from the corresponding PDE at the same final time. The two approaches produce nearly identical profiles and density levels, especially in the bulk. We observe a shift in the position of the PDE front, which is slightly more advanced than the discrete model front. The PDE model allows arbitrarily small densities to propagate, while the discrete model requires a finite number of bacteria per site to sustain forward motion, which slows the leading edge.

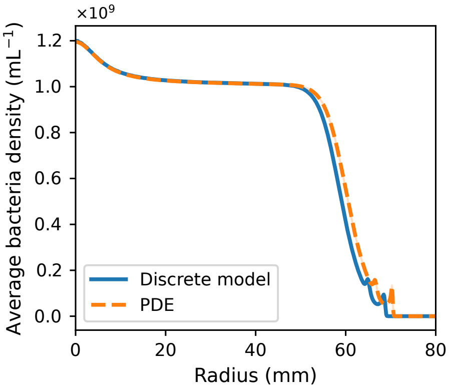


Figure S3: Radially averaged bacterial density profiles for simulations without infection. Solid line: discrete stochastic model; dashed line: deterministic PDE. Both models use identical parameters and are compared at the same final time t=19 h.

### S4 Infected Front Roughness Measure

To assess the geometric irregularity of the phage infection front we introduce a quantitative measure, allowing for comparisons between different spatial patterns to determine their degree of irregularity. The front roughness measure is defined as

$$\xi = \frac{A_{\text{out}}}{A_{\text{infected}}}, \tag{14}$$

where $A_{\text{infected}}$ is the total area of the infected surface and $A_{\text{out}}$ is the area of the infected surface lying outside a reference quarter circle of area equal to $A_{\text{infected}}$. Each simulation produces a $m$-dimensional lattice $I$, representing the number of infected bacteria in each infected state at every spatial site. We first sum over the infection-state dimension to obtain a two-dimensional lattice of total infected bacteria per site. This lattice is then binarized using a threshold $I \geq I_{\text{thr}}$, where the threshold depends on the lattice depth as $I_{\text{thr}} = 100 \cdot \Delta z$ (Figure S4, left). From this binary mask, we construct a solid infected region by filling the background connected to the initial site $(0, 0)$ using 4-connectivity, ensuring that the infected domain forms a single connected region. The total infected area $A_{\text{infected}}$ is computed as the number of lattice sites in this solid region.
To define a smooth reference shape, we construct a quarter circle with the same area $A_{\text{infected}}$. Its radius is determined analytically as $r = \sqrt{\frac{4A_{\text{infected}}}{\pi}}$, and a corresponding binary quarter-circle mask is generated on the same lattice grid. This reference shape is overlaid with the infected region (Figure S4, center), and

the area $A_{\text{out}}$ is computed as the number of infected sites that lie outside the quarter circle (Figure S4, right).

The ratio $\xi$ therefore measures the fraction of infected area that extends beyond the quarter-circle, which acts as a smooth reference. A compact, approximately circular infection front yields $\xi \approx 0$, whereas increasingly irregular fronts with pronounced protrusions result in larger values of $\xi$. Since both $A_{\text{out}}$ and $A_{\text{infected}}$ are measured in pixel units, $\xi$ is dimensionless.

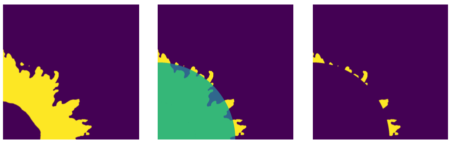

Figure S4: Computation of the infection front roughness measure $\xi$. From left to right: the infected lattice is binarized using an intensity threshold; the resulting infected region is filled to obtain a single solid domain (yellow) and overlaid with a reference quarter circle of equal area (green); finally, the infected area lying outside the quarter circle is identified and used to compute the roughness measure.

Since the computation of the front roughness measure $\xi$ requires binarizing the infected lattice using an intensity threshold $I_{\text{thr}}$, to assess the robustness of our results we performed a sensitivity analysis by varying the threshold over a range $I_{\text{thr}} \in [100, 900]$, while keeping all other parameters fixed.

The upper value $I_{\text{thr}} = 900$ corresponds to the largest threshold for which the infected region forms a compact, connected surface across all parameter combinations considered in this study. Conversely, the lower value $I_{\text{thr}} = 100$ represents a threshold that includes sites with relatively small numbers of infected bacteria, providing an estimate of the infected region extent. The intermediate value $I_{\text{thr}} = 500$ is used throughout the main text.

Figure S5 shows the front roughness $\xi$ as a function of the product of adsorption rate and burst size $\eta \cdot \beta$ for three threshold values. While the absolute magnitude of $\xi$ varies with the threshold, the qualitative behaviour remains unchanged. In particular, a pronounced peak in front roughness is observed at intermediate values of $\eta \cdot \beta$ for all thresholds considered. This indicates that the emergence of a rough infection front is not an artifact of the specific threshold choice, but reflects a robust feature of the underlying infection dynamics. Based on this analysis, we conclude that the threshold $I_{\text{thr}} = 500$ provides a reliable compromise between capturing a well-defined infected front and avoiding excessive sensitivity to low-density fluctuations, and we therefore adopt it as the default value in the main text.

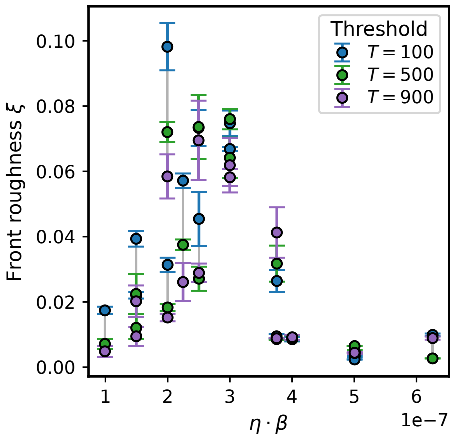


Figure S5: Sensitivity of the front roughness measure $\xi$ to the infection threshold. Front roughness is shown as a function of $\eta \cdot \beta$ for three different binarization thresholds, $I_{\text{thr}} = 100$, 500, and 900. The qualitative behaviour is preserved: in all cases, a peak in front roughness is observed at intermediate values of $\eta \cdot \beta$. Error bars indicate the standard error of the mean over 3 independent simulation replicates.

### S5 Effects of Spatial Discretization

To examine how spatial resolution influences our results, we compared simulations performed at different lattice spacings in both the deterministic PDE model and the discrete stochastic model (DM). All simulations were run on the same physical domain (80 mm × 80 mm) using identical physical parameters, and evolved for 19 hours. Figure S6a shows results from the PDE model simulated with grid spacings of $\Delta x = 0.10$ mm and $\Delta x = 0.05$ mm. The two runs produce similar bacterial and infection fronts, confirming that the overall dynamics are robust to changes in spatial resolution. However, we can observe some geometric differences: at coarser resolution ($\Delta x = 0.10$ mm), the front appears slightly flattened along the horizontal and vertical directions, while the finer grid yields a more circular, isotropic shape. This deformation arises from lattice anisotropy intrinsic to spatial discretizations: the numerical approximation of diffusion and chemotaxis permits transport only between horizontally and vertically aligned grid points, under-representing diagonal propagation. As $\Delta x$ decreases, diagonal propagation is represented more accurately, and this anisotropy decreases. In general, the PDE model shows numerical convergence with decreasing $\Delta x$: refining the grid mainly reduces anisotropy without affecting too much the position of the front or the overall dynamics.
We next tested the effect of spatial discretization in the discrete model (DM), where bacteria and phages are represented as integer-valued populations per lattice site. Simulations were performed with $\Delta x = 0.10$ mm and $\Delta x = 0.05$

mm, while keeping all physical quantities and parameters identical. As shown in Figure S6b, both simulations produce qualitatively similar spatial patterns, with comparable front morphology, chemoattractant-driven fronts, and irregular infection structures. Nevertheless, as also observed for the PDE simulations, the positions of both the outer bacterial front and the infection front differ slightly between the two spatial resolutions, likely due to small discretization effects in the representation of diffusion and chemotaxis. Given that the qualitative behaviour is preserved, we adopted $\Delta x = 0.10$ mm for all main-text simulations, which provides a good balance between computational efficiency and spatial resolution.

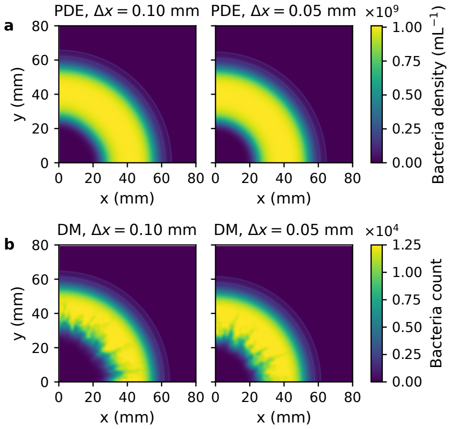


Figure S6: Effect of spatial resolution on deterministic and stochastic model results. (a) Deterministic PDE simulations at two spatial resolutions, $\Delta x = 0.10$ mm (left) and $\Delta x = 0.05$ mm (right). Both runs produce similar bacterial and infection fronts, with the coarser grid showing slight horizontal and vertical flattening due to lattice anisotropy, while the finer grid yields a more circular front. (b) Discrete stochastic model (DM) simulations at $\Delta x = 0.10$ mm (left) and $\Delta x = 0.05$ mm (right). The overall front morphology and chemoattractant-driven structures are qualitatively consistent between the two cases, although the positions of the bacterial and infection fronts differ slightly. The coarser grid was used in the main text as it captures the same qualitative behaviour while substantially reducing computational cost.

To address the concern that the use of a quarter-domain with reflective boundary conditions could introduce artificial symmetry or bias the measurement of front roughness, we performed an additional simulation using the full domain with a centrally placed inoculum. Figure S7 shows the total bacterial density at the final time in this configuration. The system develops a circular expanding bacterial front and a surrounding infection ring, with spatial irregularities qualitatively similar to those observed in the quarter-domain simulations. We can observe the same effects of spatial resolution mentioned above. We further quantified this comparison using the roughness measure described in Section S4, adapted to the full-domain geometry by defining the reference shape as a full circle of equal infected area. For the quarter-domain simulations at $\beta = 15$ and $\eta = 2.0 \times 10^{-8}$, we obtain a roughness index $\xi = 0.076 \pm 0.005$ (mean $\pm$ standard deviation over $n = 3$ independent replicates). The corresponding full-domain simulation yields $\xi = 0.066$. To compare with the quarter-domain roughness measure, we also partitioned the spatial domain into four equal quadrants and compute the front roughness measure independently in each quadrant, obtaining an average $\xi = 0.062 \pm 0.009$ (mean $\pm$ standard deviation over the four quadrants). Given the computational cost of full-domain simulations, we limited this analysis to a representative parameter set, but the results indicate that the roughness measure is not an artifact of the reduced simulation geometry.

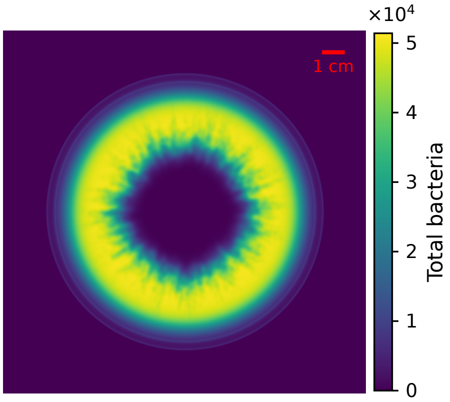


Figure S7: Full-domain simulation with central inoculation. Total bacterial density at t=19 h for a discrete stochastic model simulation performed on the full 160 mm × 160 mm domain, with bacteria and phages initially placed at the centre. Parameters are $\beta = 15$ and $\eta = 2.0 \times 10^{-8}$ mL h$^{-1}$. The system develops a circular expanding bacterial front and an infection front with irregular structure, qualitatively similar to those observed in quarter-domain simulations.

### S6 PDE simulations with inhomogeneous phage inoculum

To investigate whether spatial inhomogeneities in the initial phage inoculum alone can generate macroscopic roughness of the infection front, as proposed by Ping et al. [1], we performed two-dimensional PDE simulations with inhomogeneous initial phage distributions. In their supplementary material, Ping et al. consider one-dimensional simulations in which heterogeneity is introduced across independent realizations by varying the initial phage density, while maintaining spatial uniformity within each run, and show that the position of the phage front varies substantially between realizations. Here, we instead test the same hypothesis in a two-dimensional system.

We considered two classes of inhomogeneous phage inocula. First, we implemented a random inoculum in which phages were distributed within a circular region of radius 0.5 mm around the origin. At each lattice site within this region, the order of magnitude of the initial phage density was drawn independently from a Gaussian distribution for $\log_{10} P$, with mean $\log_{10}(P_0)$ and standard deviation 0.5, where $P_0 = 10^9$ mL$^{-1}$ denotes the initial phage density used in the homogeneous simulations. The resulting inoculum was then normalized to ensure that the total phage number matched that of the homogeneous reference case. Second, to impose a stronger and more controlled spatial heterogeneity, we constructed an explicitly inhomogeneous inoculum in which the total initial phage mass was distributed over three lattice sites: two located along the coordinate axes (sites $(0, 5)$ and $(5, 0)$) and one along the diagonal (site $(4, 4)$). The initial phage distribution is shown in Figure S8a. To conserve total phage mass, each site was assigned a density $P_0/3$. As in the previous simulations, bacteria were initialized in a single lattice site $(0, 0)$ with density $B_0$. Both inoculation protocols introduce substantial initial heterogeneity and produce the same final pattern. We focus on the latter case because it represents a more extreme and spatially structured perturbation than the random inoculum, and therefore provides a more stringent test of whether deterministic PDE dynamics alone can sustain front roughness. In contrast to the one-dimensional simula-

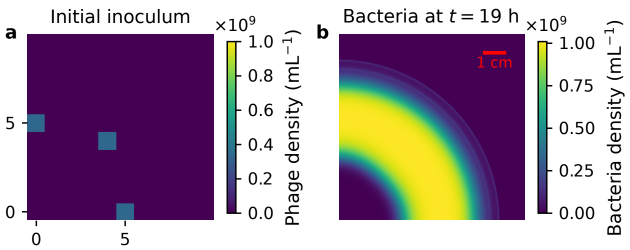


Figure S8: PDE simulation with inhomogeneous phage inoculum. (a) Zoomed view of the initial phage density field, showing the spatially inhomogeneous inoculum. (b) Total bacterial density at $t = 19$ h. Despite the initial inhomogeneity, the bacterial front remains smooth and radially symmetric.

tions of Ping et al. [1], our approach introduces heterogeneity within a single spatial realization rather than across an ensemble of independent runs. As a consequence, the initial phage inhomogeneities are subject to diffusion and adsorption dynamics in two dimensions, which rapidly smooth local variations in phage density near the inoculation region. This results in a largely uniform, radially expanding front at later times, as shown in Figure S8b. While inoculum inhomogeneity can influence early-time dynamics locally, it does not generate persistent front roughness in the deterministic two-dimensional PDE framework considered here, suggesting that additional mechanisms are required to sustain rough infection fronts.

### S7 Comparison of mean quantities with Experimental data

A direct quantitative comparison between the stochastic simulation results presented here and the experimental data of Ping et al. [1] is challenging. Front roughness is especially difficult to compare quantitatively across experiments and simulations: published experimental images are affected by imaging noise, which would make the roughness measure highly sensitive to thresholding choices. In addition, biological variability could lead to differences in pattern morphology across experimental realizations.

For these reasons, we focus on observables that are well defined in both experiments and simulations. In particular, we consider the outer radius of the bacterial front and the width of the annular bacterial region. In the work of Ping et al. [1], these quantities are reported in the form of colormaps.

To extract these observables and compare them with the results of Ping et al. [1], we performed simulations using the same initial inocula and adsorption rates. From these simulations, we analyzed the spatial distribution of total bacteria at the final simulation time. The final patterns are are shown in Fig. S10a. To quantify the spatial structure, we compute radially averaged bacterial density profiles, which capture the mean radial structure of the expanding population while averaging out angular fluctuations.

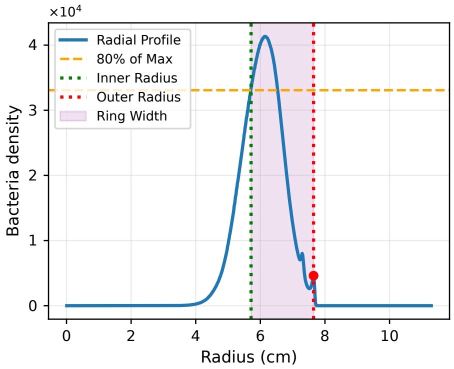


Figure S9: representation of methods to extract radial observables from simulation data. The radially averaged bacterial density profile $\langle B(r) \rangle$ is shown for a representative simulation. The outer radius $R_{\text{out}}$ is defined as the position of the outermost local maximum of the profile (red marker). The inner radius $R_{\text{in}}$ is defined as the smallest radius at which the density exceeds 80% of the peak value (green dashed line). The shaded region highlights the resulting ring width $W = R_{\text{out}} - R_{\text{in}}$.

From the radial profile $\langle B(r) \rangle$, we define the following quantities:

- **Outer radius $R_{\text{out}}$:** The outer radius is defined as the radial position of the outermost local maximum of the radially averaged bacterial density profile. We identify all local maxima of $\langle B(r) \rangle$ and select the peak at the largest radius.

- **Inner radius $R_{\text{in}}$:** The inner radius is defined as the smallest radius at which the radial density exceeds 80% of the peak density,

$$\langle B(R_{\text{in}}) \rangle \geq 0.8 \max_r \langle B(r) \rangle. \tag{15}$$

- **Ring width $W$:** The width of the annular region is then defined as

$$W = R_{\text{out}} - R_{\text{in}}. \tag{16}$$

Figure S9 illustrates this procedure for a representative simulation, showing the radial density profile together with the detected outer peak, the threshold defining the inner radius, and the resulting ring width. Using the definitions above, we computed $R_{\text{out}}$ and $W$ across the full parameter sweep in adsorption rate $\eta$ and initial phage number $P_0$. The resulting colormaps are shown in Figure S10b and Figure S10c. Increasing adsorption efficiency or initial phage inoculum leads to a systematic decrease of the outer radius, reflecting stronger phage suppression of bacterial expansion. At the same time, the width of the annular region decreases. These trends mirror those reported experimentally and in the deterministic modelling of Ping et al. [1], despite the presence of stochastic front roughness in our simulations that is absent from deterministic PDE descriptions. We note that their deterministic model does not include adsorption to already infected bacteria, whereas this process is included in our model. This difference, together with the stochastic nature of our simulations, may contribute to quantitative discrepancies between the two approaches.

### S8 Alternative Sources of Stochasticity in Lysis Dynamics

In the Erlang implementation of the infection cycle, the mean latency is fixed at $1/\psi(n)$, while the coefficient of variation decreases as $1/\sqrt{m}$. Thus, increasing the number of infection states narrows the distribution of lysis times and suppresses stochasticity, yielding smoother fronts. To understand why this happens, and which parts of the distribution are more important for roughness, we considered mixtures of subpopulations with different lysis rates. Upon infection, bacteria are assigned with small probability $p_J$ to a subpopulation of infected bacteria with a different lysis rate $\psi_J$, while the majority $(1 - p_J)$ enters the main subpopulation with lysis rate $\psi_I$. Both subpopulations proceed through

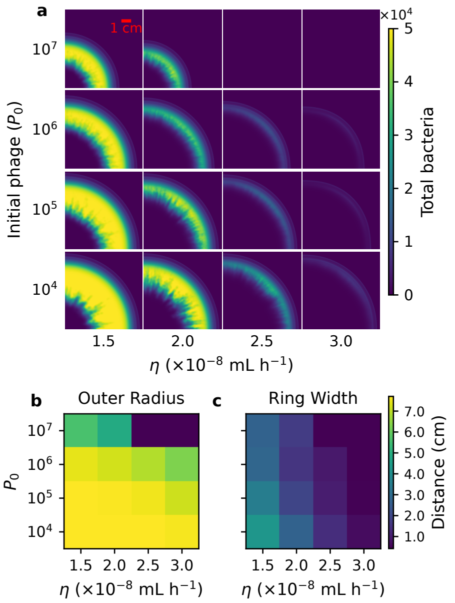


Figure S10: Comparison of simulated bacterial patterns with radial observables across adsorption rate and initial phage inoculum. (a) Final spatial distributions of total bacteria for varying adsorption rate $\eta$ and initial phage number $P_0$. (b) Outer radius $R_{\mathrm{out}}$ extracted as the position of the outermost peak in the radially averaged bacterial density profiles for each parameter combination. (c) Corresponding ring width $W = R_{\mathrm{out}} - R_{\mathrm{in}}$, where $R_{\mathrm{in}}$ is defined using an 80% threshold of the maximum radial density.

3 steps of infection and release the same burst size at lysis. This generates an effective latency distribution

$$f(\tau) = (1 - p_J)\,\mathrm{Erlang}(m, \psi_I)(\tau) \; + \; p_J\,\mathrm{Erlang}(m, \psi_J)(\tau), \tag{17}$$

which may be shifted toward shorter times (if $\psi_J > \psi_I$) or extended toward longer times (if $\psi_J < \psi_I$), as shown in Figure 6a. From the simulations shown in Figure 6b we can observe that adding a minority of fast-lysing infections restores roughness more effectively than adding a minority of slow-lysing ones. This indicates that rare early lysis events are the key driver of rough infection fronts, while longer latency outliers add variability in timing but do not strongly affect front geometry. The mixture model shows that the suppression of roughness at high $m$ arises mainly from the loss of the early-lysis tail in the latency distribution.

To further test whether front roughness could be recovered by increasing infection encounters, we varied the adsorption rate $\eta$ while keeping fixed $m = 5$. As shown in Figure S11, higher $\eta$ values shift the infection front outward—reflecting faster overall infection spread—but do not restore the sharper branching characteristic of low-$m$ systems. This confirms that stochasticity in lysis timing, not encounter frequency, is the dominant factor driving the emergence of rough fronts.

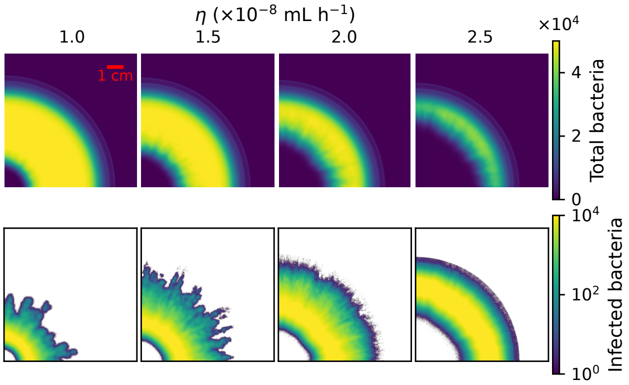


Figure S11: Infection patterns at 19 hours for $m = 5$ at varying adsorption rates. Infection patterns are shown for $\eta = 1.0$–$2.5 \times 10^{-8}$ mL h$^{-1}$. Increased $\eta$ shifts the infection front outward but does not recover the branches seen at lower $m$.

In addition to variability in lysis rates, we also explored heterogeneity in the swimming behaviour of infected cells. We used the two subpopulations construction from above: upon adsorption, cells enter the main infected class $I$ with probability $(1 - p_J)$ or the minority class $J$ with probability $p_J$. Here, all lysis parameters are fixed across subpopulations, so the latency distribution $f(\tau)$ is unchanged relative to the pure Erlang case. The difference is in the chemotactic coefficients: the minority class responds more weakly or more strongly to gradients

$$\chi_J = \alpha\,\chi_I, \qquad \alpha < 1 \text{ (less responsive)} \quad \text{or} \quad \alpha > 1 \text{ (more responsive)}. \tag{18}$$

Introducing either less responsive ($\alpha = 0.8$) or more responsive ($\alpha = 1.2$) infected cells broadens the distribution of lysis positions relative to the homogeneous case, even though the latency distribution itself remains narrow. However, the consequences for front roughness are asymmetric, as shown in Figure S12. Less responsive cells tend to lyse slightly behind or around the bulk front and therefore have little visible effect on its shape. By contrast, more responsive cells can travel further up the gradient and lyse ahead of the bulk population, producing rare forward outliers that act as new seeding events. It is these forward outliers — whether generated by shorter latency times or by enhanced motility — that are responsible for restoring roughness to the infection front.

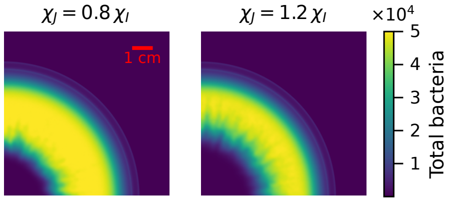


Figure S12: Spatial infection fronts under heterogeneity in swimming behaviour of infected cells. Snapshots at 19 hours show total bacterial density when the subpopulation of infected cells $J$ have reduced motility ($\chi_J = 0.8 \cdot \chi_I$, left) or enhanced motility ($\chi_J = 1.2 \cdot \chi_I$, right) compared to the subpopulation of infected cells $I$.
